# Sir2 non-autonomously controls differentiation of germline cells in the ovary of *Drosophila melanogaster*

**DOI:** 10.1101/631176

**Authors:** Champakali Ayyub, Ullas Kolthur-Seetharam

**Affiliations:** 710, Bhaskara, TIFR Colony, Homi Bhabha Road, Mumbai 400005, India; Department of Biological Sciences, Tata Institute of Fundamental Research, Mumbai 400005, India

**Author notes:** Corresponding Author: Champakali Ayyub.

**Keywords:** *Sir2*, Escort cell, *Drosophila*, Dpp, Germline stem cell

## Abstract

In *Drosophila* ovary, germline stem cells (GSCs) reside in a somatic cell niche that provides them signals necessary for their survival and development. Escort cells (ECs), one of the constituents of the niche, help in differentiation of GSC daughter cells. Since nutritional state is known to affect oogenesis, we set out to address the role of a metabolic sensor. NAD-dependent Sir2 is known to acts as a regulator of organismal life-span in a diet dependent manner. Our current study reveals that Sir2 in somatic cells is necessary for germline differentiation. Specifically, Sir2 in ECs upregulates Dpp signalling giving rise to tumorous germaria. In addition to this non-autonomous role of Sir2 in regulation of the germline cell homeostasis, we have demonstrated that EC-specific Sir2 has a role in attributing the identity of Cap cells as well as in de-differentiation of germline cells. Our study also shows that a genetic interaction between *Sir2* and *upd2* is important for the development of germline cells. Thus, we provide novel insights into the role of Sir2 in ovary development.

## Introduction

Stem cells need signals from their specialized environment or ‘niche’ for their survival and development. An established system to study this stem cell-niche interaction is the ovary of *Drosophila melanogaster* that contains 14-16 tubular structures called ovarioles. Anteriormost part of the ovariole is called germarium that harbours germline stem cells (GSCs). The ovarian niche is made up of somatic cells namely Terminal Filament cells (TFCs), Cap cells (CpCs) and Escort Cells (ECs) (Decotto and Spradling, 2005; Xie and Spradling, 1998). In the niche, several signalling pathways (*e.g.*, BMP, JAK/STAT, Wnt and Hh) act on the germline cells in a coordinated manner to regulate their maintenance, proliferation and differentiation (Decotto and Spradling, 2005; Kai and Spradling, 2003; Liu et al., 2015; Lopez-Onieva et al., 2008; Luo et al., 2015; Wang et al., 2008; Xie and Spradling, 1998, 2000). However, cell autonomous factors that modulate these non-autonomous signaling pathways, are less understood. Moreover, since oogenesis is known to be affected by dietary inputs, if and how metabolic sensors affect this developmental process needs to be unraveled.

Bone Morphogenetic Protein like molecule BMP homologue Decapentaplegic (Dpp) is known to be a major signalling pathway. Dpp derived from the CpCs act directly on the GSCs to suppress a differentiating factor encoded by *bag of marbles* (*bam*) and, thus, helping the GSCs for their self-renewal. However, this signal gets attenuated sharply causing daughter cells of GSCs (called Cystoblast or CB) situated just one cell diameter away to undergo differentiation by expressing Bam. This process is followed by four incomplete divisions of the CB to form a sixteen cell cyst that moves towards the posterior end and gets wrapped up by somatic cells, eventually giving rise to an ovum (Chen and McKearin, 2003c; McKearin and Ohlstein, 1995; McKearin and Spradling, 1990; Song et al., 2004). Anteriorly, CBs are tightly wrapped by extensions from the ECs and are eventually encapsulated by the somatic cells as the cyst moves towards the posterior end. Interestingly, this physical interaction between the CB and the ECs is necessary to trigger the process of CB differentiation for the formation of the cysts (Banisch et al., 2017; Gancz et al., 2011; Kirilly et al., 2011; Liu et al., 2015; Maimon et al., 2014).

The conserved JAK/STAT pathway in *Drosophila* consists of three different ligands (Unpaired: Upd1, Upd2 and Upd3), a trans-membrane receptor called Domeless, a Janus Tyrosine Kinase Hopskotch and a transcription factor Stat92e. This pathway is activated upon binding of ligands to Dom that triggers Hopskotch to phosphorylate Stat92e; phosphorylated Stat92e moves to the nucleus to regulate expression of its downstream genes (Hombria and Brown, 2002). This is one of the pathways important for the communication between the soma and germline cells in the ovary (Decotto and Spradling, 2005; Lopez-Onieva et al., 2008; Wang et al., 2008).

Stat92E is highly expressed in ECs and mutants of Stat92E show a loss of GSC phenotype. On the other hand, upregulation of Upd in ECs gives rise to ectopic GSC-like cells in the germaria, which together indicate that the JAK/STAT signaling in ECs is important for the GSC maintenance (Lopez-Onieva et al., 2008; Song et al., 2004; Xie and Spradling, 2000). Originally ECs were shown to provide structural support to the germarium. Subsequently, ECs were described as a source of low levels of Dpp and now are known to regulate the extent of Dpp signalling in the niche. ECs are also responsible for both GSC self-renewal and differentiation of the CBs. Interestingly, ECs produce low levels of Dpp when the JAK/STAT signalling gets activated in response to Upd originated from the CpCs. This EC-derived Dpp is stated to be responsible for the maintenance of a pool of partially differentiated germline cells, which can dedifferentiate to GSCs (Decotto and Spradling, 2005; Eliazer et al., 2011; Kirilly et al., 2011; Liu et al., 2015; Wang et al., 2008).

Interestingly, it is not only the signals originating in somatic cells comprising the niche sheltering the GSCs that prompt them to proliferate. Nutritional state or stress too provide essential cues for maintenance of the GSCs. Significant loss in number of GSCs has been reported in females shifted from rich to poor diet or with age. Thus, studies on interaction between nutritional state of the organism and maintenance of GSCs in the niche have established that, in addition to the intrinsic and extrinsic signals from their niche, Insulin signalling (IIS) too plays an important role in *Drosophila* ovary. IIS regulates the number of CpCs and also the CpCs-GSC adhesion. Thus, IIS controls the size of the niche as well as the mechanism for GSC maintenance in a Dpp-independent manner (Drummond-Barbosa and Spradling, 2001; Gancz and Gilboa, 2013; Hsu and Drummond-Barbosa, 2009). Further, recent studies have emphasised the importance of both IIS and Target of rapamycin (TOR) pathways during development of ovarian stem cells and their niche (Gancz and Gilboa, 2013).

Sirtuins, a family of NAD^+^-dependent deacylases, are known to couple the metabolic status to various cellular processes. SIRT1, whose orthologues are conserved from bacteria to mammals, is a key regulator of transcription in the nucleus. The *Drosophila* homologue *Sir2* plays a major role in glucose and fat metabolism as well as in insulin signaling in a tissue specific manner with effects on organismal survival. Reports have highlighted the ability of Sir2 to exert tissue non-autonomous control in several contexts. Specifically, fat body *Sir2* establishes tissue communication with muscles and the insulin producing cells (via JAK/STAT signaling), which are important for energy homeostasis (Banerjee et al., 2012b, 2013; Bishop and Guarente, 2007; Helfand and Rogina, 2003; Palu and Thummel, 2016; Reis et al., 2010; Rogina and Helfand, 2004). Further, Sir2 is known to affect organismal lifespan in a diet-dependent manner (Banerjee et al., 2012a; Rogina and Helfand, 2004; Tissenbaum and Guarente, 2001).

Since diet has been shown to impinge on oogenesis, a possible role of Sir2, specifically in a non-autonomous manner, has not been elucidated. In wild type germaria, Sir2 expression has been observed in germline cells but is confined only to regions 1 and 3 with its level going down in regions 2a and 2b. This observation is concomitant to the fact that Sir2 has a significant function in the regulation of pachytene checkpoint and checkpoint is suppressed in regions 2a and 2b. Failure to suppress pachytene checkpoint, as has been found in loss-of-function mutations of *Sir2*, gives rise to defects in oocyte determination (Joyce and McKim, 2010; Pek et al., 2012). In this study, we have reported that Sir2 is also present in somatic cells and our findings underscore a non-cell autonomous control of the *Sir2* in *Drosophila* ovary. We establish that loss of *Sir2* in somatic cells affects differentiation of germline cells and *Sir2* epistatically interacts with both Dpp and Upd2 to exert its control over oogenesis.

## 2. Materials and methods

### 2.1. Fly Stocks

The following stocks were used: *w*^*1118*^ was considered a control genotype. The Gal4 stock *c587-Gal4* and *Sir2-GFP* (CB02821) were obtained from M. Buszczak (University of Texas, Southwestern Medical Centre, Dallas). All other Gal4 stocks, mutant of *Sir2* (*Sir2*^2A-7-11^), *P[bamP-GFP], dpp*RNAi A (BL 31531), *dpp*RNAi B (BL 31530) and two mutants of *dpp* (BL 36528: *dpp*^*hr56*^*cn*^*1*^*bw*^*1*^*/SM6a* and BL 2069: *dpp*^*hr92*^*cn*^*1*^*bw*^*1*^*/SM6a*), *upd2*RNAi (BL33949) were obtained from the Bloomington Stock Centre. Sir2RNAi (CG5216:23201/GD) was obtained from the Vienna Drosophila RNAi Center (VDRC). Flies were grown and maintained at 25°C on standard *Drosophila* medium unless otherwise mentioned. For Gal4-driven crosses, flies were grown and aged at 29 °C.

### 2.2. Immunostaining and antibodies used

Immunostaining of ovaries was performed using a method described earlier (Ayyub et al., 2015). Briefly, ovaries were dissected in PBS and fixed in 4% Formaldehyde for 20 minutes. After washing in PTX (PBS with 0.3% Triton X-100), the ovaries were incubated in primary antibody diluted in PTX overnight at 4°C. Next day, they were thoroughly washed in PTX and then incubated in secondary antibody for 2 hours at room temperature. After another course of washes, the ovaries were incubated in 1 µg/ml of Hoechst for 10 minutes followed by an additional set of washes and mounting in Vectashield.

Primary antibodies were used in this study at the following dilutions: rat anti-Vasa (1:25), mouse anti-Spectrin (3A9, 1:40), mouse anti-β-galactosidase (1:100), mouse anti-Armadillo (N2 7A1, 1:50), mouse anti-Lamin C (LC28.26, 1:30) from the Developmental Studies Hybridoma Bank, Rabbit anti-phosphohistone H3 (1:500) from Sigma, rabbit anti-Caspase 3 (1:250) from Santa Cruz Biotechnology, rabbit anti-pERK from Cell Signaling Technology (1:100), rabbit anti-pMad (1: 200) and rabbit anti-GFP (1:10,000) from Abcam, guinea pig anti-Traffic jam (1:10,000) was a gift from D. Godt, University of Toronto, Canada. All the secondary antibodies used at 1:300 dilution (Alexa Fluor-488, -568, -647 conjugated goat anti-mouse, -rat or -rabbit antibodies) and rabbit anti-GFP (1:10000) were obtained from the Molecular Probes, Bangalore, India and were.

Olympus FV1000 confocal microscope was used to acquire images. Images were processed by Adobe Photoshop 7.0 and Olympus Fluoview version FV10-ASW4.0 viewer.

### 2.3 fertility assay

Single virgin females were crossed to 3-4 control males to set up lines. Progeny from each line were counted.

### 2.4 dedifferentiation experiment with hs-bam construct

We followed the method described in (Liu et al., 2015). Briefly, flies grown at 18°C and aged for 4-5 days at 25°C were heat shocked at 37°C for 1h 20 mins, followed by a recovery period of 2h 20 mins and another heat shock at 37 °C for 1h 20 mins. Flies were then maintained on fresh food at 25°C till immunostaing.

### 2.5 Statistical analysis

We have presented the data as mean ± standard deviation and P-values were calculated using one-way ANOVA in OriginPro 2015. For drawing graphs, Microsoft Excel was used.

## 3. RESULTS

### 3.1. Escort cells of the germarium express Sir2

Earlier studies showed that in the *Drosophila* ovary, Sir2 is expressed in germline cells (Pek et al., 2012), however, if it is expressed in somatic cells is still unknown. We used *Sir2-GFP*, an enhancer trap line where GFP expression is controlled by the *Sir2* locus (Buszczak et al., 2007; Kelso et al., 2004; Reis et al., 2010), to study Sir2 expression patterns in somatic cells of the ovary. As identified by their characteristic triangular shape and location in the anterior part, ECs in regions 1 and 2a of the germarium were found to express Sir2 (Fig. 1B-B’). Considering the importance of ECs in maintenance of GSCs and differentiation of CBs (Decotto and Spradling, 2005; Eliazer et al., 2011; Liu et al., 2015; Song et al., 2004), this EC-specific expression of Sir2 led us to investigate the effects of EC-specific down-regulation of *Sir2* on germline cells of *Drosophila* ovary.

**Fig. 1.**
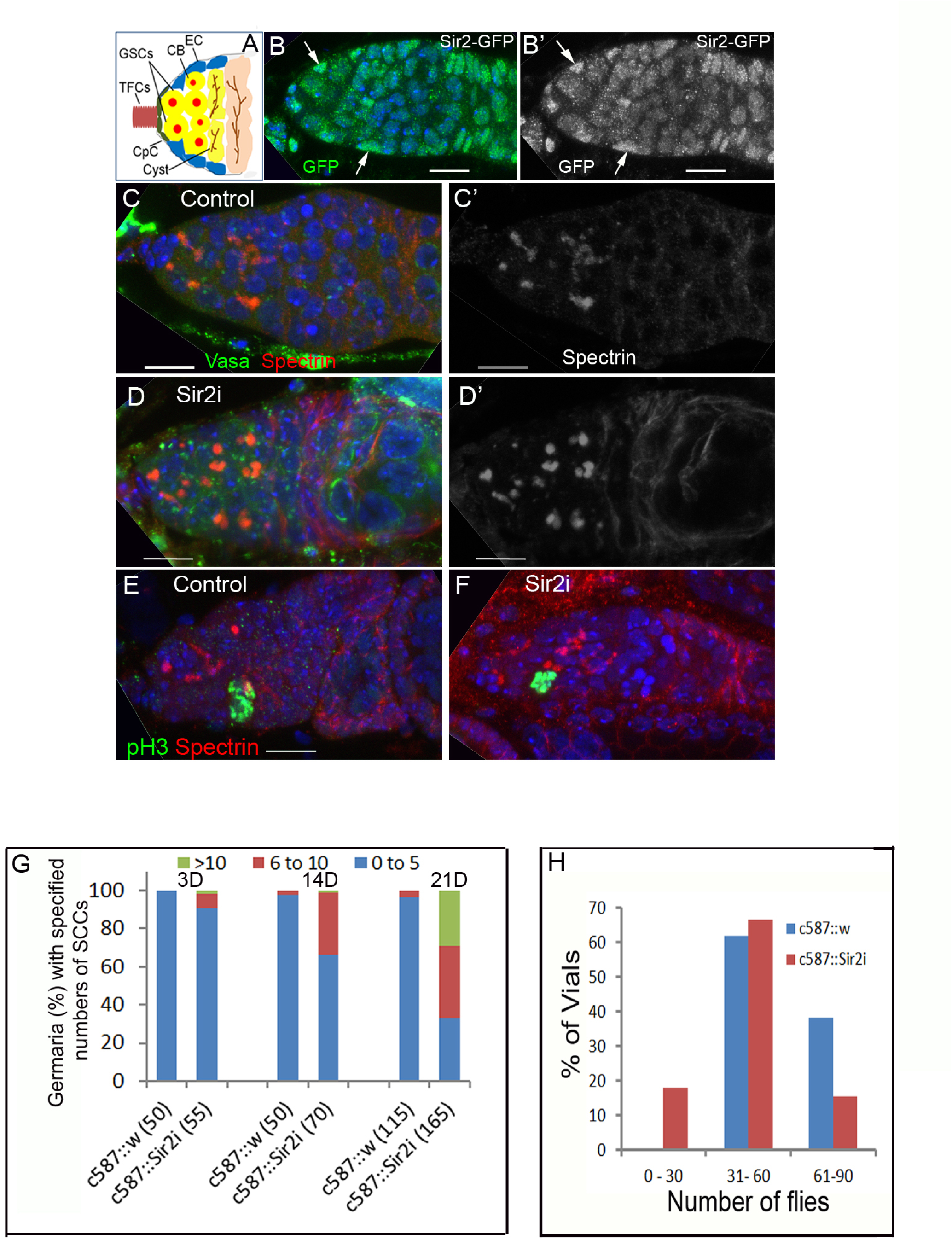
EC-specific Sir2 is required in the development of germline cells. (A) A schematic diagram of the anterior part of the *Drosophila* germarium where different cell types are as follows: escort cells (ECs), cystoblasts (CBs), germline stem cells (GSCs), terminal filament cells (TFCs) and cap cells (CpCs). The spherical Spectrosomes present in both GSCs and CBs branch into Fusomes in cysts. (B-B’). Expression of *Drosophila* Sir2 in germarium as observed in an enhancer-trap line where GFP expression is under the control of the *Sir2* locus. Prominent expression of Sir2 is present in the ECs (marked by white arrows) (C-D) Germaria from 21D old ovaries of control (C-C’) and Sir2i (D-D’) immunostained for Vasa (green: germline cell marker) and Spectrin (red) that marks the Spectrosomes and Fusomes. EC-specific down-regulation of Sir2 causes accumulation of SCCs (D-D’). (E-F) Immunostaining of germaria from control (E) and *Sir2i* (F) ovaries with phosphorylated histone H3 (green) shows that the ectopic cells in *Sir2i* continue to divide similar to the control. (B-F) DNA marker Hoechst is in blue. Scale bar: 10µm. (G) Quantification of germaria with the indicated number of SCCs at three different ages. Numbers in parenthesis indicate the number of germaria scored. (H) Fertility of 21D old females of genotype *c587::Sir2RNAi* decline with respect to the control.

### 3.2. Reduction of Sir2 in ECs produces germaria with ectopic GSC-like cells

To investigate whether Sir2 in the ECs had any role in ovary development and oogenesis, we used the Gal4-UAS system (Brand and Perrimon, 1993). The *c587-Gal4* line that expresses in ECs and early follicle cells in adult flies (Song et al., 2004; Yang et al., 2017) (Fig. S1A) was used to drive *Sir2*-RNAi transgene to down-regulate *Sir2* only in the ECs. Both GSCs and CBs contain α-Spectrin-rich spherical structure called Spectrosome, which in differentiating cells branches through the cytoplasmic bridges of the cyst and are known as Fusomes (Lin and Spradling, 1995, 1997; Lin et al., 1994). Thus, the presence of Fusomes in a germarium is a signature of differentiation (de Cuevas and Spradling, 1998; Lin et al., 1994). Generally, in a wild-type germarium, 2-5 Spectrosome containing cells are present and earlier studies have shown that the number of GSCs do not alter significantly over a period of 3 weeks (Panchal et al., 2017). Interestingly, downregulation of *Sir2* in ECs resulted in a progressive phenotype. Specifically, while germaria from 3D old females from *c587-Gal4::Sir2i* looked normal, its weak tumorous phenotype at 14D progressed to a tumorous phenotype by 21D as scored by the accumulation of extra Spectrosome-containing cells (SCCs; Fig. 1G). In 21D old control females (*c587-Gal4::w*), there were 2-5 Spectrosome-carrying cells in 96.5% and 6-10 GSC-like cells in 3.6% germaria (n=120 germaria). However, in *c587-Gal4::Sir2i* ovaries of the same age, 32.9% were normal, 37.7 % with 6-10 SCCs per germarium and 29.4% were mildly tumorous (Figs. 1C-D & S1B-C). Accumulation of these extra SCCs in germaria arising from EC-specific depletion of *Sir2* indicates a possible impairment in differentiation of germline cells. Staining with phosphorylated histone H3 revealed that the ectopic cells in *c587-Gal4::Sir2i* continue to divide similar to the control (Figs. 1E-F). However, on assaying for activated Caspase 3, a marker for programmed cell death, we found increased apoptosis in *c587-Gal4::Sir2i* (Figs. S3A-B). As expected, fertility of 21D old *c587-Gal4::Sir2i* females was significantly reduced with respect to the control females (Fig. 1H).

In a parallel experiment, we also employed *tj-Gal4* that expresses in both ECs and CpCs (Li et al., 2003) to knockdown *Sir2* simultaneously in both of these cell types. Here, we observed an age-dependent germarial phenotype that was similar to but weaker than the only EC-specific abrogation of *Sir2* (Fig. S2A). Incidentally, fertility of these females tested 21D after eclosion in single pair crosses was also substantially reduced in comparison with the control (Fig. S2B). Given this, we restricted our further investigations into the role of *Sir2* in oognesis to its knockdown only in ECs *i.e. c587-Gal4::Sir2i* (henceforth denoted only as *Sir2i*).

### 3.3 EC-specific Sir2 restricts Dpp signalling in the niche and controls differentiation of CBs

Accumulation of GSC-like cells in germaria occurs when there is an increase in Dpp signalling or a down-regulation of Bam in germ cells (McKearin and Ohlstein, 1995; Song et al., 2004). Hence, we were prompted to determine if *Sir2i* enhances Dpp signaling in the ECs. We assayed this by studying the expressions of two of the important markers of the pathway namely activated Mad and Dad. While, Mothers against Dpp (Mad) is required for Dpp signal transduction, Dad or Daughters against Dpp, which is downstream to Dpp acts as a negative regulator of the pathway. In wild-type germarium, expression of pMad (phosphorylated Mad) is restricted only to the GSCs where Dpp level is high. However, Dad as scored using an enhancer trap Dad-lacZ reporter line (where lacZ expression is under the control of Dad), which is expressed in both GSCs and the early CBs (Tsuneizumi et al., 1997). As reported, the wild-type germaria had prominent pMad expression only in GSCs (2.27 ± 0.83, n=62) occupying the anterior-most region of the germarium (Fig. 2A). In contrast, in *Sir2i* germaria, the pMad-expressing cells (3.02 ± 0.92, n=82; P<0.0001) were located even several cell diameters away from the CpCs (Figs. 2B and SD-D’). As mentioned in earlier studies (Liu et al., 2015), pMad expression in the SCCs outside the niche is weak when compared to that in the GSCs, suggesting a decrease in the Dpp level. Interestingly, the number of GSCs in *Sir2i*, identified with respect to their position in the germaria (1.80 ± 0.59) was comparable to that of the control (1.81 ± 0.62; P=1). This indicated that some of the pMad positive single cells are GSC-like in *Sir2i* germaria, but are located away from the niche. Moreover, in comparison with the control germaria (4.98 ± 1.9, n=55; P<0.0001; Figs. 2C-C’), germaria with EC-specific *Sir2* knockdown also exhibited higher number of Dad-lacZ-expressing cells (9.76 ± 3.5, n=70; Figs. 2D-D’), some of which are displaced from the niche. Thus, our results from Dad-lacZ-expression clearly illustrate that an absence of *Sir2* in the ECs generate a population of SCCs with some ectopic GSC-like cells as well as an accumulation of early CBs. Importantly, these suggest that Sir2 expression in ECs regulates Dpp signalling and that reduction of *Sir2* causes an ectopic increase in Dpp, which results in defective germline development.

**Fig. 2.**
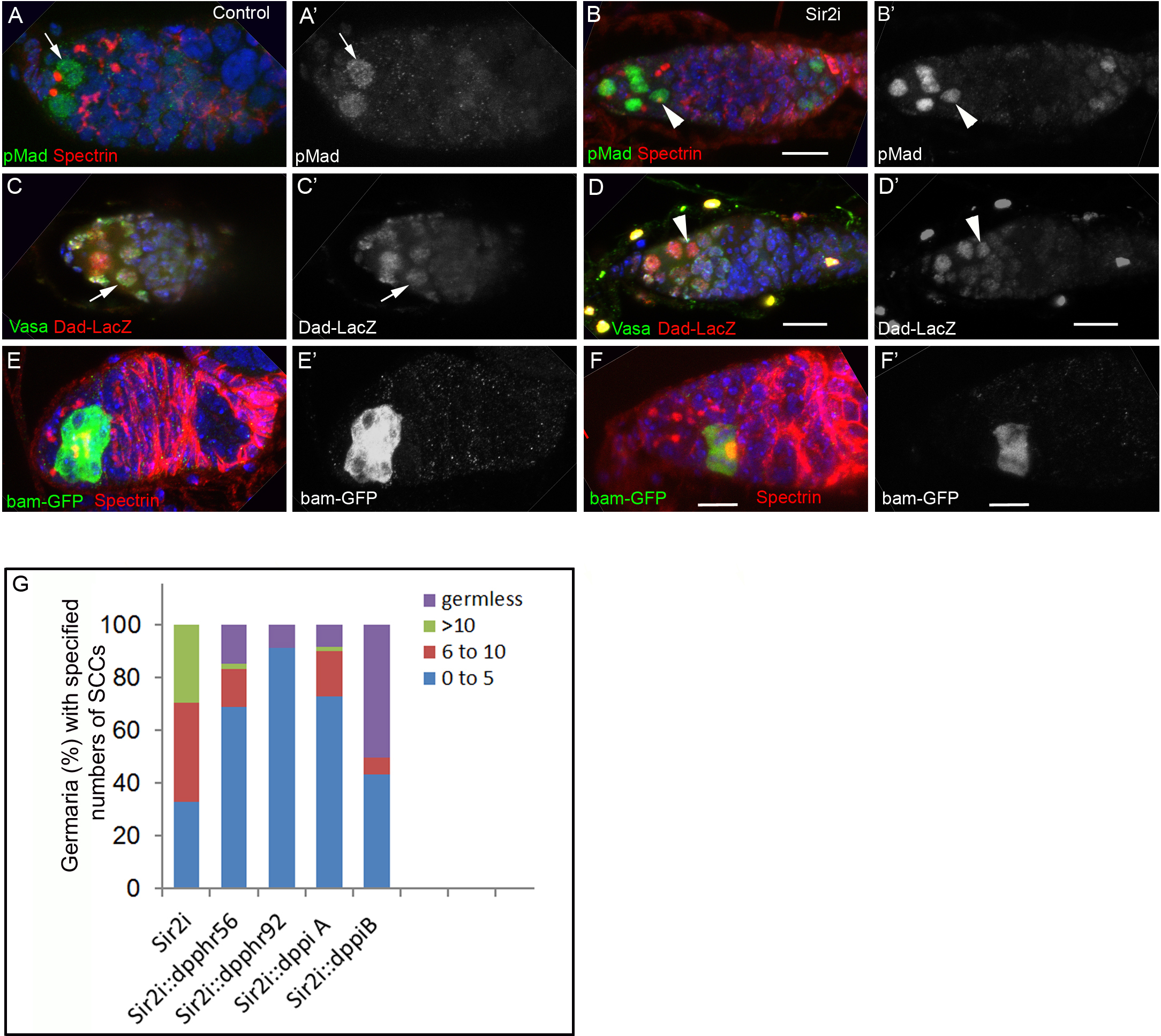
EC-specific knockdown of *Sir2* causes an extension of Dpp signalling. (A-B) A control germarium has pMad (green in A and B) expression in two GSCs at the anteriormost region (white arrow) (A-A’) whereas in the germarium of *Sir2i*, it expresses even away from the niche (though at a lower intensity; white arrowhead) showing that the ectopic cells are GSC-like cells (B-B’). (C-D) In control germaria, prominent expression of Dad-lacZ (red in C and D) is observed only in GSCs whereas in CBs its expression is weak (white arrow) (C-C’). In *Sir2i*, however, level of Dad-lacZ expression in CBs (based on their location away from the niche: white arrowhead) is comparable to that in the GSCs (D-D’). (E-F) GFP expression (green in E and F) in control germarium carrying *bamP-GFP* (E-E’) is strong in the dividing cyst (E) but in the germarium of *Sir2i*::*bamP-GFP* with ectopic SCCs, it expresses weakly (F-F’). (A-F) DNA marker Hoechst is in blue. Scale bar: 10µm. (G) Quantification of germaria with the indicated number of SCCs where n is at least 50 germaria for each genotype except in *Sir2i* (n=165) and *Sir2i::dppiB* (n=122).

While Dpp dependent repression of Bam in GSCs inhibits differentiation, down-regulation of Dpp in cystoblasts causes de-repression of Bam and hence initiates differentiation (Chen and McKearin, 2003b; Kai and Spradling, 2003). Therefore, we were interested to verify whether the ectopic expression of Dpp suppressed *bam* expression giving rise to the germarial phenotype in *Sir2i* females. Towards this, we used *P[bamP-GFP]* that carries a *bam* promoter driven GFP-reporter, which is expressed only in the differentiating cysts (Chen and McKearin, 2003a). The expression pattern of GFP in germline cells is typical by being absent in GSCs, low in CBs and highly expressed in dividing cysts. Presence of *bam*-positive germline cells reduced from 94% in control (n=50) to 25% in the experimental (n=65) germaria (Fig. 2E-F), as reproted. Among the *Sir2i* germaria with *bam*-positive germline cells, the intensity of GFP is reduced compared to the control. In some germaria with excess SCCs, single cells express GFP at a low intensity showing that they are early CBs (Fig. S1F). These observations suggest that upregulation of Dpp in *Sir2i* is sufficient to suppress Bam expression in the germline cells and consequently leading to an increase in the number of SCCs in the germaria.

Given that EC-specific depletion of *Sir2* caused excess Dpp and a concomitant accumulation of extra SCCs, we anticipated that a simultaneous reduction of Sir2 and Dpp in the ECs would rescue the tumorous phenotype of germaria. Our data with two different *dpp* RNAi transgenes (denoted by A and B) in the ECs and two *dpp* mutants (*dpp*^*hr56*^ and *dpp*^*hr92*^ present in one copy) revealed that there indeed was a partial rescue of the tumorous phenotype and consequently more germaria with Fusome-carrying cysts were obtained in comparison with *Sir2i* (Fig. 2G). These results indicate that *Sir2* expression in EC has a role to restrict Dpp beyond the niche. Interestingly, for the *dpp* RNAiB transgene, in addition to the rescue to the normal phenotype, there was a large number of empty germaria (50%, n=122 germaria), which is a characteristic phenotype of the loss of *dpp* (Fig. S4). Together, these findings clearly illustrate that Sir2 functions upstream of Dpp signalling.

### 3.4 Downregulation of Sir2 in ECs does not change the number of CpCs but confers a mixed identity of the CpCs

Our data presented so far reveal that the up-regulation of Dpp signaling in *Sir2i* resulted in excess SCCs. CpCs, located at the tip of the germarium, being the major source of Dpp form a key component of the niche and are crucial for the maintenance of GSCs (Xie and Spradling, 1998, 2000). Interestingly, ECs in the niche restrain the extent of Dpp signaling and thus support the differentiation of CBs (Xie and Spradling, 2000). Anteriormost part of the germarium harbours 5-7 CpCs and formation of the CpCs is governed by Notch signalling that acts at the pupal stage to determine the number of CpCs (Song et al., 2007). As expected, ectopic Notch signalling increases the number of CpCs. Presence of extra CpCs causes an extension of the Dpp signalling beyond the niche that eventually gives rise to tumorous germaria (Song et al., 2004; Song et al., 2002; Xie and Spradling, 1998, 2000). We wanted to check whether an increase in the number of CpCs was responsible for the rise in the Dpp signalling. By immunostaining with a marker called Armadillo, we found that with respect to the control (6.7± 1.47, n=50) there was no significant increment (P=0.08) in the number of CpCs in *Sir2i* (5.9 ± 1.16, n=60; Fig. 3A-B). These results suggest that the tumorous phenotype in *Sir2i* is possibly not because of an elevated Notch signalling.

**Fig. 3.**
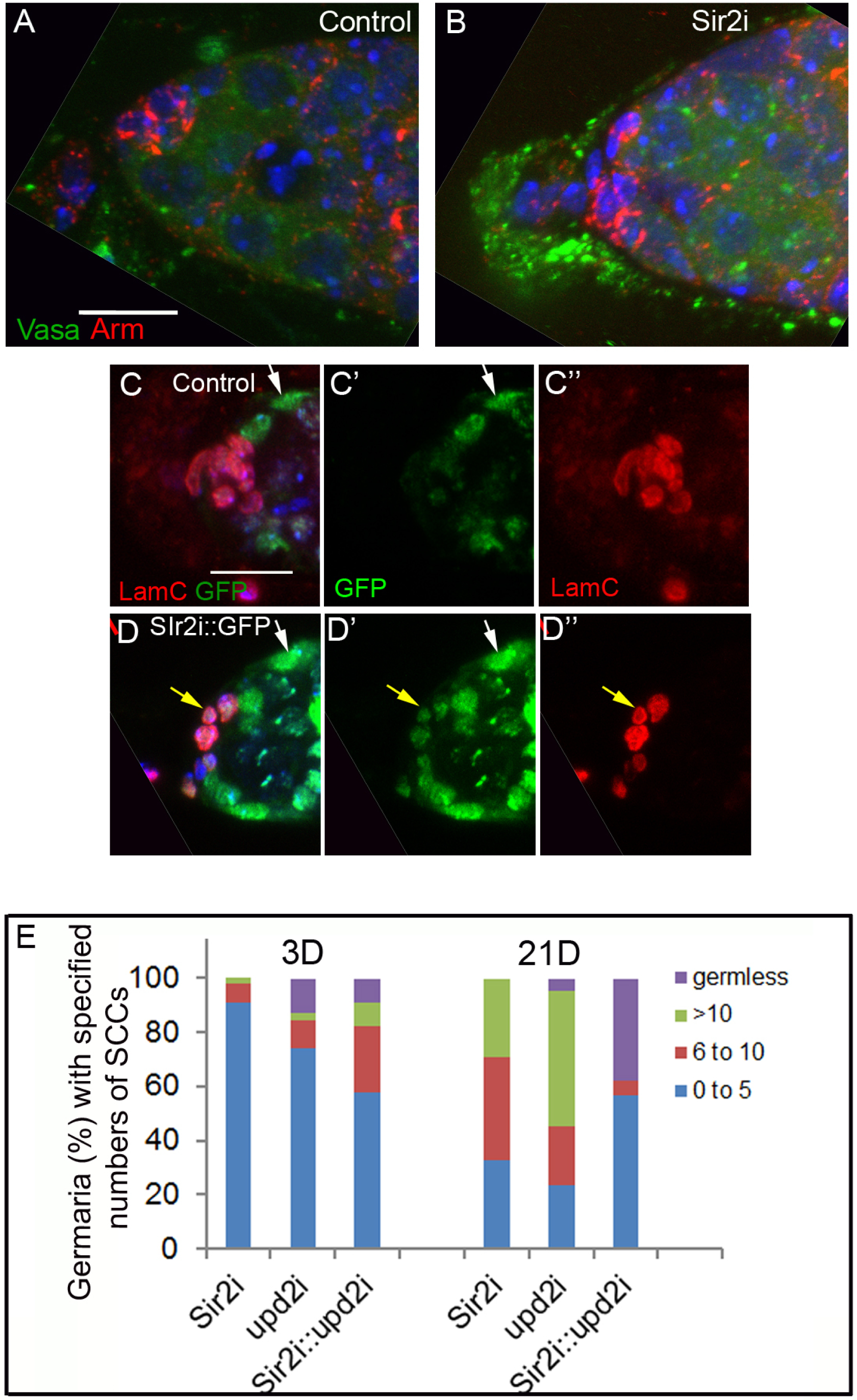
Downregulation of *Sir2* in ECs does not change the Cap cell number but gives rise to CpCs with a mixed identity. (A-B) Cap cells marked by Armadillo (red) are similar in number in control (A) and *Sir2i* (B) germaria. Germline marker Vasa is in green. (C-D) In control c587::UAS-GFP (C-C”) and *Sir2i::GFP* (D-D”) ECs were marked by GFP (green) as shown by white arrows and CpCs and TFCs were marked by Lamin C (red). In *Sir2i::GFP*, CpCs too express Lamin C (yellow arrow). (A-D) DNA marker Hoechst is in blue. Scale bar: 10µm. (E) Quantification of germaria with the indicated number of SCCs obtained from 3D and 21D old ovaries from *Sir2i::upd2i* along with the controls. (n=at least 40 germaria for 3D old females and n=at least 90 germaria for 21D females).

Next, we explored the other possibility that there was a change in cell fate among the somatic cells in the niche. In the germarium, nuclear envelope of CpCs and TFCs express Lamin C, and is used as a marker. As expected, in ovaries of *c587-Gal4::UAS-GFP*, the GFP expression and Lamin C staining were exclusive to ECs and CpCs/TFCs, respectively (Fig. 3C-C”). Interestingly, in *c587-Gal4::Sir2i::UAS-GFP*, while Lamin C expression was normal, GFP expression was observed in ECs as well as in CpCs and faintly in TFCs (Fig. 3D-D”). These data suggest that depletion of *Sir2* in ECs have altered the identity of the CpCs, which have characteristics of both ECs and CpCs leaving the properties of ECs unaltered. This finding is unlike the fate-changes described earlier *viz* with the loss of Lsd1 (a histone demethylase) where CpCs and ECs exhibit markers of both cell types (Eliazer et al., 2011) or with EC-specific down-regulation of histone H1 where ECs attain the likeness of CpCs (Yang et al., 2017). So, our results suggest that *Sir2* has a role in cell fate determination of CpCs.

Another reason to have an up-regulated Dpp is an increase in number of somatic cells in the germarium. We used anti-Traffic jam antibody that marks somatic cells in the ovary (Li et al., 2003) and found that in the germarium, their number in *Sir2i* is comparable with the corresponding control (Fig. S3C-D). Additionally, an interaction between the niche and the germline cells, which is essential for the latter to proliferate, is mediated by the EGFR-MAPK pathway (Gilboa and Lehmann, 2006; Guo and Wang, 2009; Hayashi et al., 2009; Liu et al., 2010). From pERK staining, a marker of MAPK pathway, we found that there was no significant change in *Sir2i* with respect to the control (Fig. S3E-F).

### 3.5 EC-specific Sir2 interacts with Upd2 to restrain Dpp signalling

As mentioned in the introduction, a feedback regulation between Dpp and JAK/STAT signaling plays an important role in ovary development (Decotto and Spradling, 2005; Lopez-Onieva et al., 2008; Wang et al., 2008). Importantly, changes in both these signaling mechanisms are required for the maintenance of GSCs and function in the same direction. Interestingly, JAK/STAT controls Dpp signalling in ECs at a low level to maintain GSCs in a partially differentiated state (Liu et al., 2015) (Lopez-Onieva et al., 2008; Wang et al., 2008; Xie and Spradling, 1998). Similar to Dpp, ectopic expression and reduction of Upd in somatic cells cause accumulation of extra SCCs and empty germaria, respectively.

We have previously shown that an interaction between Sir2 and Upd2 maintains organismal metabolic homeostasis, wherein Sir2 acts as an upstream regulator of Upd2 (Banerjee et al., 2017). Thus, we wanted to explore whether there was a similar interaction between Sir2 and Upd2 in ovaries as well. Towards this, we compared phenotypes of 3D and 21D old ovaries of the genotypes *Sir2i*, c587:: *upd2-RNAi* (hereafter referred to as *upd2i*) and the double knock-down in ECs (hereafter referred to as *Sir2i::upd2i*). In young flies, *Sir2i* ovaries lacked any perceivable phenotype (Fig. 3E). However, consistent with earlier studies, in *upd2i* 12.8% (n=40) germaria were germless and they also showed a mild tumorous phenotype (10.2% of germaria with 6-10 SCCs and 2.5% with >10 SCCs). Interestingly, *Sir2i::upd2i* displayed germless phenotype in 8.8% germaria (n=45) but there arose a strong tumorous phenotype in 24.4% germaria with 6-10 SCCs and 8.8% with >10 SCCs. As a result, in germaria of the double-knockdown ovaries, occurrence of normal germaria went down to 57.7%, which was much lower than those observed in individual knockdowns (90.9% in *Sir2i* and 74.3% in *upd2i*). Thus, this set of data shows that there is an interaction between *Sir2* and *upd2* in the ovaries at young age, where Sir2 seems to act upstream of Upd2.

As already mentioned, by 21D *Sir2i* develops tumorous germaria. At the same age, the germless phenotype exhibited by *upd2i* was weaker (4.6% germaria, n=90) than that of the young flies. But notably, these flies exhibited a strong tumorous phenotype with 22.1% germaria having 6-10 SCCs and 50% germaria having >10 SCCs. Interestingly, this unique phenotype in *upd2i* was ameliorated when it was in combination with *Sir2i.* There were more normal-looking germaria (56.6%, n=90) compared to the individual genotypes whereas the germless phenotype (37.8% germaria) became predominant. These further corroborated the earlier findings and supported that there is an interaction between *Sir2* and *upd2*, and indicated their importance in germline differentiation.

### 3.6 De-differentiation of the germline cells is dependent on Sir2 in ECs

Previous studies have demonstrated that under special circumstances cysts undergoing differentiation are capable of reprogramming themselves to generate GSCs. A positive regulator of this process has been found to be a low level of EC-specific Dpp that maintains germline cells in a partially differentiated state (Kai and Spradling, 2004; Liu et al., 2015). Since we found that the EC-specific down-regulation of Sir2 caused a moderate up-regulation of Dpp beyond the niche, we wanted to determine whether that affected de-differentiation. Differentiation can be induced by an ectopic expression of Bam in GSCs (Chen and McKearin, 2003a; Ohlstein and McKearin, 1997). Thus, we used a *hs-bam* transgene in the background of *c587-Gal4* as well as *c587-Gal4::Sir2-RNAi*, along with a relevant control. In these three genotypes, we assayed the number of GSCs using the markers pMad one day (1D) and seven days (7D) after heat-shock (AHS) treatment to induce *bam* expression (Liu et al., 2015). In control germaria, there were two to five pMad positive cells present at both the time points (n=70 germaria). However, only 9.2% germaria (n=70) had pMad positive cells because of the Bam-induced differentiation of GSCs in *c587-Gal4::hs-bam* 1D AHS. Interestingly, at 7D AHS, the number of germaria with GSCs increased up to 70.0% (n=60) clearly indicating de-differentiation of the artificially differentiated cysts that was observed at 1D AHS (Fig. 4E)).

**Fig. 4.**
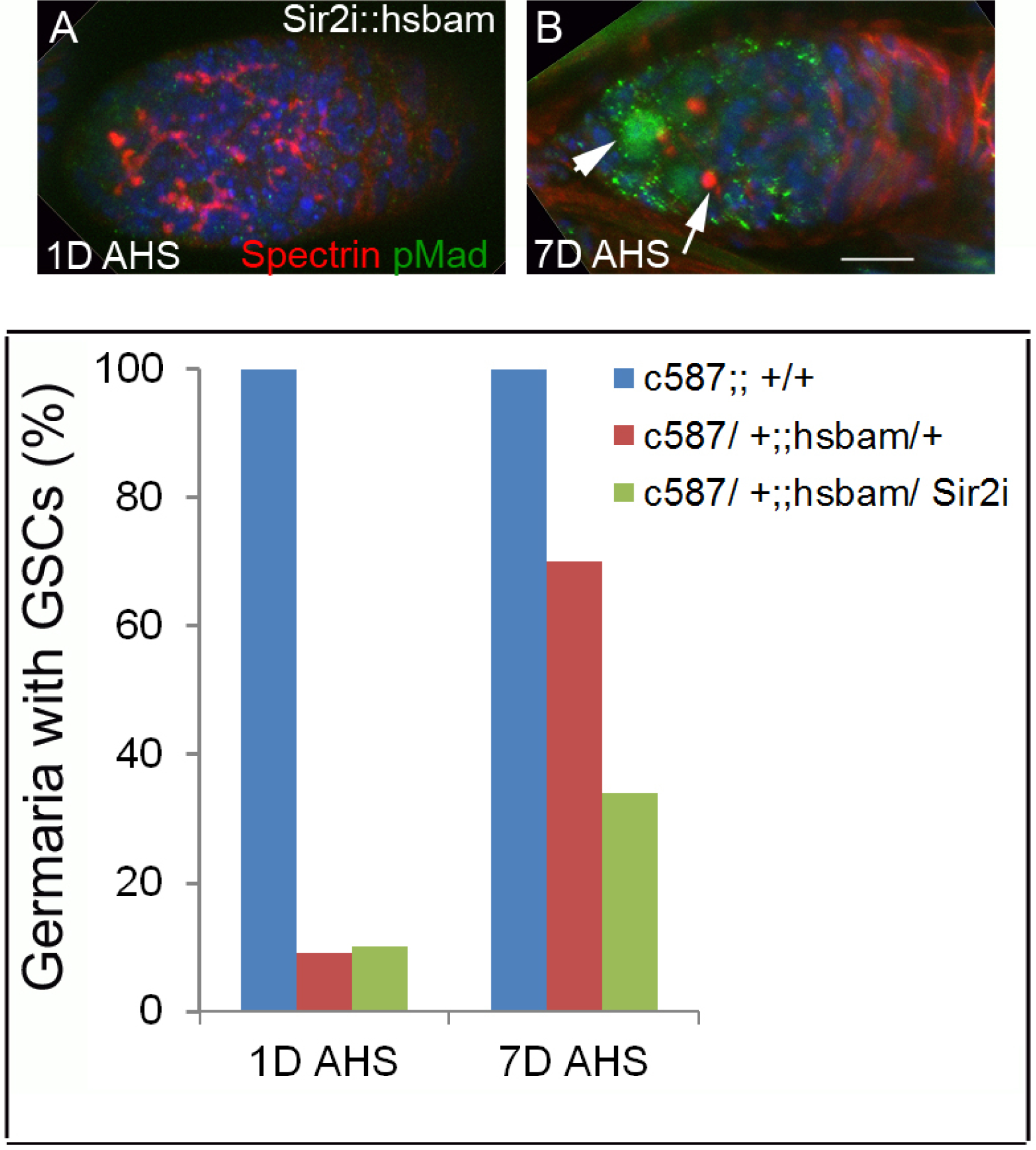
Reduction of Sir2 in ECs affects dedifferentiation in the germaria. (A-B) Most germaria of *c587::Sir2-RNAi::hs bam* 1D AHS does not have any pMad-positive GSCs but have only Fusome-carrying cysts (pMad is in green and Spectrin is in red) (A) but 7D AHS, a fraction of the germaria scored, had pMad-positive GSCs (white arrowhead) and Spectrosomes (white arrow) (B). DNA marker Hoechst is in blue. Scale bar: 10µm. (C) Quantification of germaria with GSCs in control and two other genotypes with *hs-bam* in the background at 3D and 7D AHS.

Similar to *c587-Gal4::hs-bam, c587-Gal4::Sir2i::hs-bam* was found to have only 10.1% of germaria (n = 85) with GSCs in their niche 1D AHS. However, at 7D AHS, there were 33.9% (n = 60) of the germaria with pMad-positive cells, which was much less than the *hs-bam* control. These data suggest that here too some of the Bam-induced Fusome-carrying cells have possibly undergone de-differentiation to replenish GSCs in the niche. However, it is important to note that EC-specific abrogation of *Sir2* dampened the de-differentiation potential of differentiating cysts into GSCs. In other words, our findings imply that in addition to the low level of Dpp in ECs (Liu et al., 2015), *Sir2* has a significant role to play in the de-differentiation of ectopic cysts to GSCs in the germaria. However, how exactly Sir2 acts in de-differentiation in a Dpp-independent manner remains to be elucidated.

## DISCUSSIONS

Regulation of germ-cell niche and differentiation programmes have been shown to be dependent upon spatiotemporal cues involving orchestrated gene expression. In the context of ovaries, down-regulation of several histone-modifying enzymes (*viz, Lsd1, Set1* and H3K4 tri-methylase) and histone H1 itself in ECs have been shown to impinge on the Dpp pathway giving rise to Spectrosome-carrying tumours (Eliazer et al., 2011; Wang et al., 2011; Xuan et al., 2013; Yang et al., 2017). In this study, we have unravelled a novel role for NAD-dependent deacylase Sir2, a well-known regulator of transcription and chromatin remodelling, in determining differentiation during oogenesis. Specifically, we highlight the importance of Sir2 in the ECs in controlling germline-fates in a non-autonomous manner.

Several publications on EC-specific overexpression of Upd have shown an interaction between JAK/STAT and the Dpp pathways (Decotto and Spradling, 2005; Kai and Spradling, 2003; Wang et al., 2008). Studies on other signaling mechanisms have also highlighted the importance of the non-autonomous role of ECs in regulating the number of germline cells (Eliazer et al., 2011; Kirilly et al., 2011; Liu et al., 2010; Wang et al., 2011). However, the relevance of a metabolic sensor in somatic cells exerting control over germline cells is poorly understood. Sir2 is known for its roles in metabolic homeostasis and aging (Banerjee et al., 2012a; Banerjee et al., 2012b, 2013; Banerjee et al., 2017; Burnett et al., 2011; Delaney et al., 2011; Kaeberlein et al., 1999; Rogina and Helfand, 2004). In this context, we have shown that EC-specific abrogation of Sir2 causes accumulation of SCCs in an age-dependent manner. Based on our results from immunostaining and genetic studies, we have demonstrated that the tumorous phenotype arises because of an up-regulation of Dpp.

Earlier reports have illustrated that egg production in *Drosophila* depends on diet and further studies demonstrated that insulin signalling in combination with the signals emanating from the niche govern GSC proliferation (Drummond-Barbosa and Spradling, 2001; Hsu and Drummond-Barbosa, 2009; Hsu et al., 2008; LaFever and Drummond-Barbosa, 2005). However, the importance of upstream regulators is hitherto unknown. Upd2 is now regarded as a key factor in mediating inter-tissue communication between the fat-body and insulin producing cells in the brain (Rajan and Perrimon, 2012). We have earlier shown that the NAD-dependent Sir2 in fat-body exerts long range effects on insulin signaling via Upd2 and fat homeostasis. Importantly, Sir2 is epistatic to Upd2 in regulating insulin secretion from a distance (Banerjee et al., 2012b, 2013; Banerjee et al., 2017). During ovary development, earlier studies elucidated the independent roles of Upd2 and Sir2. Therefore, this report not only reiterates the importance of these molecules in ovary development but also establishes an interplay between them in maintaining germline differentiation. Specifically, the data presented in section 3.5 (Fig. 3E) reveal that for the two time points considered, simultaneous down-regulation of *Sir2* and *upd2* yielded different phenotypes. Percentage of germaria displaying wild type phenotype remained the same in *Sir2i::upd2i* in both 3D and 21D old flies. However, a comparison of tumorous and germless phenotypes reveals that these are completely different in 3D old and 21D old ovaries. Therefore, our findings indicate that *Sir2* and *upd2* interact in an age-dependent manner to regulate the Dpp signalling essential for the maintenance of GSCs. Moreover, we demonstrate that EC-specific Sir2 has multiple roles including its non-autonomous function in maintaining homeostasis of germline cells, in de-differentiation of the partially differentiated germline cells and also in the determination of CpCs. Given this, it would be interesting to explore, in the future, the interplay between dietary inputs and Sir2 functions in orchestrating ovary development.

Aging and metabolic inputs have been shown to significantly contribute towards germ-line differentiation. However, genetic factors that control aging and their contributions emerging from the somatic cells have not been investigated thus far. In this regard, we have identified Sir2, a metabolic sensor and an anti-aging factor, as one of the key determinants of germline homeostasis in a Dpp dependent manner. Importantly, we have discovered that Sir2 in the somatic cells (ECs) is necessary for regulating germcell-niche interactions.

## Figure legends

Fig. S1. Expression pattern of *c587-Gal4* and phenotypes of *Sir2i*. (A) The Gal4-driver *c587-Gal4* expresses prominently in the ECs (white arrows). (B-C) Germaria of *Sir2i* have extra SCCs (C) compared to the control (B). (D-D’) In *Sir2i*, the pMad positive cells are also present away from the CpCs (white arrow) showing that some of the SCCs are GSC-like. (E-F) Compared to the GFP in control germaria (E), ectopic SCCs in *Sir2i* have weak GFP showing that they are early CBs (F). (A-F) DNA marker Hoechst is in blue. Scale bar: 10µm.

Fig. S2. Simultaneous knockdown of Sir2 in CpCs and ECs mildly affects the development of germline cells. (A) Quantification of germaria with the indicated number of SCCs in 21D old *tj-Gal4*-driven control and *Sir2i*. (B) Fertility of the 21D old females of *tj-Gal4::Sir2i* declines with respect to the control.

Fig. S3. EC-specific reduction of Sir2 upregulates programmed cell death but there is no appreciable change in the number of total somatic cells or in EGFR-MAPK signalling. (A-B) In comparison with the control (A), there is an elevated expression of the activated Caspase 3 (green) in *Sir2i* (B). (C-D) Immunostaining with anti-traffic jam antibody reveals that the EC-specific down regulation of *Sir2* (D) does not change the number of somatic cells with respect to the control (C). (E-F) pERK expression in *Sir2i* (F) is comparable to its expression in the control (E). DNA marker Hoechst is in blue. Scale bar: 10µm.

Fig. S4. Quantification of germaria with the indicated number of SCCs. Reduction of Dpp in ECs in *Sir2i* with *dpp* RNAiB transgene (depicted as *dppiB*), produces 43.4% of normal germaria but a large number of empty germaria (50%, n=122 germaria) too which is a characteristic phenotype of downregulation of *dpp* alone in ECs with *c587Gal4* (n=50 and n=165 germaria for *c587::dppiB* and *Sir2i*, respectively).

## Acknowledgements

We thank Prof K. Ray for his constant support and Sonam Chaudhary for helping us in the fertility assays. We also thank the Bloomington Stock Centre, the Developmental Studies Hybridoma Bank (DSHB) and the Vienna Drosophila RNAi Centre (VDRC) for the reagents. This work has been supported by a grant from TIFR-DAE (the Govt. of India).

