## Supplementary figures and images for "Sir2 non-autonomously controls differentiation of germline cells in the ovary of *Drosophila melanogaster*"

### Supplementary figures

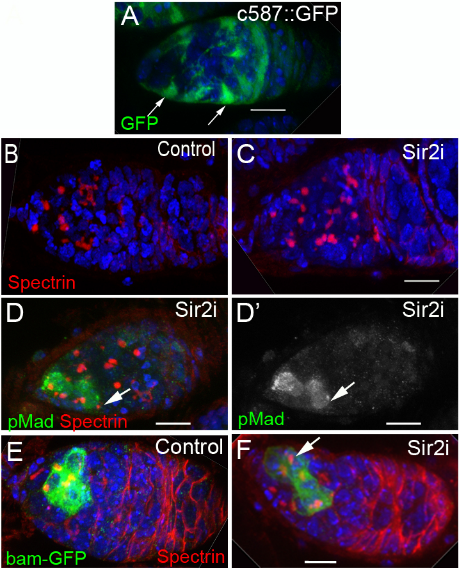

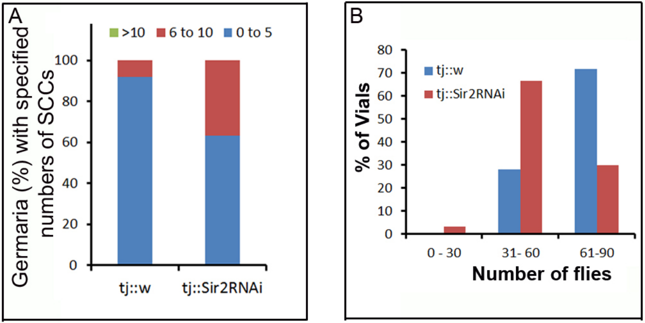

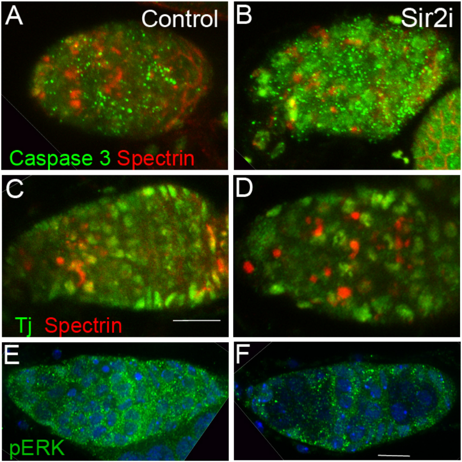


Figure S3

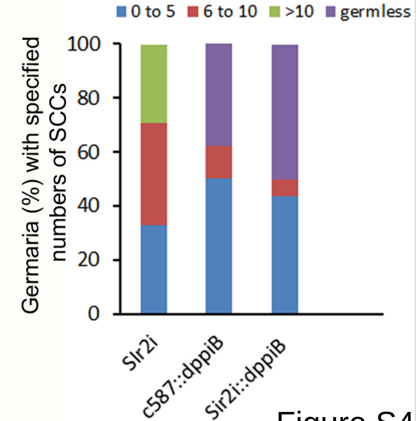
